# Ligand-binding/transcriptional repressor domain-deficient REV-ERBβ inhibits dendrite and spine formation of newborn adult hippocampal neurons

**DOI:** 10.64898/2026.08.29.746924

**Authors:** Koji Shimozaki

## Abstract

REV-ERBβ is a transcriptional repressor of nuclear receptors that regulates the circadian rhythm and plays an important role in regulation of the proliferation, differentiation, and maturation of neurons. Dysregulation of the circadian rhythm has been associated with neuropsychiatric disorders, and activation of REV-ERBs can induce anxiolytic behavior in mice. Furthermore, hippocampal neurogenesis is important in the effects of antidepressants. However, the role of REV-ERBβ in adult hippocampal neurogenesis *in vivo* at the single-cell level is not known. In this study, protein localization of REV-ERBβ in the subgranular zone of the hippocampal dentate gyrus (DG) was mainly shown in NeuN-positive neurons, and the effect of expressing a dominant negative form of REV-ERBβ lacking the C-terminal region on newborn neurons in the hippocampal DG of adult mice was examined to investigate the role of REV-ERBβ in neurogenesis. A retroviral vector containing the dominant negative REV-ERBβ or a control vector was injected into the mouse DG. At 4 weeks after injection, the morphology of dendrites and dendritic spines of newborn neurons labeled by the virus was examined. Expression of the dominant negative form of REV-ERBβ inhibited dendrite outgrowth and branching and decreased dendritic spine formation in newborn neurons in the adult mouse hippocampal DG. This study revealed a new role for REV-ERBβ in adult hippocampal neurogenesis at the single cell level, and the results will provide insight into neurogenesis in the adult brain and its relationship with psychiatric disorders.

## Introduction

The brain serves as a decision-making organ. In many mammals, including humans, new neurons are added to complex neural networks in certain regions of the brain during the lifetime of the individual. New neurons in the adult brain are generated exclusively from quiescent and active neural stem cells (NSCs) and neural progenitor cells (NPCs) in the subventricular zone (SVZ) of the ventricles and the subgranular zone (SGZ) of the hippocampal dentate gyrus (Altman and Das 1965; Eriksson et al. 1998; Gage 2000; Cameron and McKay 2001;

Alvarez-Buylla and Lim 2004; Ehninger and Kempermann 2008; Pilz et al. 2018). NSCs in the SVZ and SGZ have characteristics of astrocyte-like cells, expressing glial fibrillary acidic protein, as well as the intermediate filament protein Nestin and the SRY-box-containing transcription factor Sox2, both of which are stem cell markers. In the SGZ, type-1 NSCs, which have a radial morphology, produce type-2a NSCs that exhibit proliferative activity. These cells further differentiate into type-2b cells, which have NPC properties. After these cells differentiate into migratory neuroblasts (type-3), cell cycle arrest occurs, resulting in the formation of granule cells (Kempermann et al. 2004). In the SGZ of the adult brain, the efficiency of neurogenesis of NSCs is approximately 70%, and these cells can exhibit both self-renewal and pluripotency (Zhao et al. 2006). Most newborn neurons in the adult brain undergo survival selection during the first few days of life. Surviving newborn neurons are assumed to be functionally integrated into neural circuits of the central nervous system and contribute to brain functions such as olfaction and hippocampal-dependent learning and memory (Aimone et al. 2006; Sierra et al. 2010; Ming and Song 2011; Akers et al. 2014).

The hippocampal volume has been reported to be reduced and its function is impaired in depressed patients (MacQueen et al. 2003). However, neurogenesis in the adult hippocampus is considered to be essential for antidepressant effects and hippocampal stress tolerance (Shors et al. 2001; Zhao et al. 2008; Aimone et al. 2011; Akers et al. 2014; Johnston et al. 2016; Lehmann et al. 2013; Planchez et al. 2020). Stress inhibits the entire process of differentiation and maturation of neurons from NSCs in the hippocampus. Acute or chronic stress inhibits NSC proliferation and maturation, and this phenomenon is often observed in animal models of chronic stress-induced depression (Willner 2017; Patel et al. 2019; Cameron and Glover 2015; Lepousez et al. 2015; Sailor et al. 2017). Furthermore, the relationship between antidepressants and neurogenesis highlights a newly described biophysiological effect. Hippocampal NSCs/NPCs of rats treated with the antidepressant fluoxetine show no effect after several days of acute administration. However, chronic treatment for 2 to 4 weeks promotes hippocampal NSC/NPC proliferation and increases the number of newborn neurons (Malberg et al. 2000). In addition, chronic fluoxetine treatment has been reported to promote dendritic development and maturation of newborn neurons in a stage-specific manner (Amellem et al. 2017). These effects of antidepressants on neurogenesis and maturation are similar to the time course of the therapeutic effects in clinical practice, suggesting that antidepressant and anxiolytic treatment may increase NSC proliferation and the differentiation and maturation of newborn neurons in the adult brain (Micheli et al. 2018).

Dysregulation of circadian rhythms has also been reported to cause neuropsychiatric disorders such as depression and anxiety disorders (McClung 2007; Franken and Dijk 2009; McClung 2013; Jagannath et al. 2013; Banerjee et al. 2014). Circadian rhythm regulators are deeply involved in the regulation of NSCs and neurogenesis in the adult brain. In the neuronal differentiation mechanism of cultured adult NSCs, REV-ERB beta (REV-ERBβ, Rev-erbAβ, or NR1D2) plays an important role in regulation of proliferation, differentiation, and maturation of newborn neurons (Shimozaki 2018). In addition, a synthetic ligand that acts as an agonist to activate endogenous REV-ERBs induced wakefulness and anxiolytic behavior in wild-type mice. REV-ERBβ is an REV-ERB nuclear receptor family member that contributes to regulation of the molecular clock by directly repressing the expression of clock genes such as muscle ARNT-like protein 1 (BMAL1) and the circadian rhythm factor CLOCK. The BMAL1/CLOCK heterodimer creates a feedback loop that comprises the central mechanism of the circadian rhythm (Ko and Takahashi 2006; Kojetin and Burris 2014). Interestingly, analysis using *Rev-erbβ* gene-deficient mice and synthetic agonists suggested that wakefulness and anxiolytic behavior could be specifically regulated via REV-ERBβ (Banerjee et al. 2014). However, the molecular mechanism of REV-ERBβ-mediated anxiolysis remains unclear.

REV-ERBβ is a transcriptional repressor of nuclear receptors. It binds to the transcriptional regulatory region of the target gene at a DNA binding site in the center of the molecule and regulates transcription through its C-terminal region, which consists of a ligand binding domain and a transcriptional repression domain. In this study, a retroviral expression system consisting of REV-ERBβ lacking the C-terminal region was used as a dominant negative form to investigate the intrinsic role of REV-ERBβ in neurogenesis in the adult mouse brain. An *in vivo* investigation of adult hippocampal neurogenesis was performed using forced expression analysis of NSCs/NPCs in the hippocampal dentate gyrus of adult mice. The dominant negative form of REV-ERBβ lacking the C-terminal region inhibited dendrite outgrowth and dendritic spine formation in newborn neurons. These results may provide clues to the molecular mechanisms of REV-ERBβ-mediated anxiolytic effects.

## Results

### REV-ERBβ protein was mainly localized in the nuclei of neurons in the hippocampal dentate gyrus of the adult mouse brain

Newborn neurons in the adult brain are produced from NSCs in restricted regions such as the SVZ and SGZ. The REV-ERBβ protein is located in the nuclei and cytoplasm of cultured adult mouse and rat brain-derived NSCs, and higher levels of REV-ERBβ protein are detected in the nuclei of neurons that differentiate from cultured NSCs (Shimozaki 2018, 2021). To clarify the function of REV-ERBβ in adult neurogenesis *in vivo*, the protein expression and localization of REV-ERBβ were investigated with a particular focus on the hippocampal SGZ of the adult mouse brain. First, the hippocampal dentate gyrus of the adult mouse brain was immunostained with a combination of antibodies against REV-ERBβ, including the neuron-specific nuclear protein neuronal nuclei (NeuN) and doublecortin (DCX) (Fig. 1a). NeuN, a marker of differentiated neurons, was localized in the nucleus and was detected in the granule cell layer (GCL). The subgranular layer (SGL) between the GCL and hilus (HL) showed immunostaining signals for type 2b and type 3 neural progenitors and for DCX, a protein marker of immature newborn neurons. REV-ERBβ was mainly expressed in GCL nuclei, and most REV-ERBβ-positive cells were NeuN-positive. Although REV-ERBβ expression was also detected in some NeuN-positive and DCX-positive newborn neurons near the boundary between the GCL and SGL, the majority of NeuN-negative and DCX-positive cells were REV-ERBβ-negative. Next, the hippocampal dentate gyrus of adult mouse brains was immunostained with anti-REV-ERBβ antibodies and antibodies to the NSC/NPC cell marker Sox2 and proliferation marker Ki67 (Fig. 1b). NSCs/NPCs in the SGL are Sox2-positive, and the self-renewing and proliferating population is Ki67-positive. A small number of Sox2-positive/Ki67-negative cells were detected among the REV-ERBβ-positive cells, but the majority of REV-ERBβ-positive cells were negative for both Sox2 and Ki67. These *in vivo* results qualitatively demonstrated that the REV-ERBβ protein is mainly expressed in NeuN-positive neurons in the SGZ of the hippocampal dentate gyrus.

**Fig. 1.**
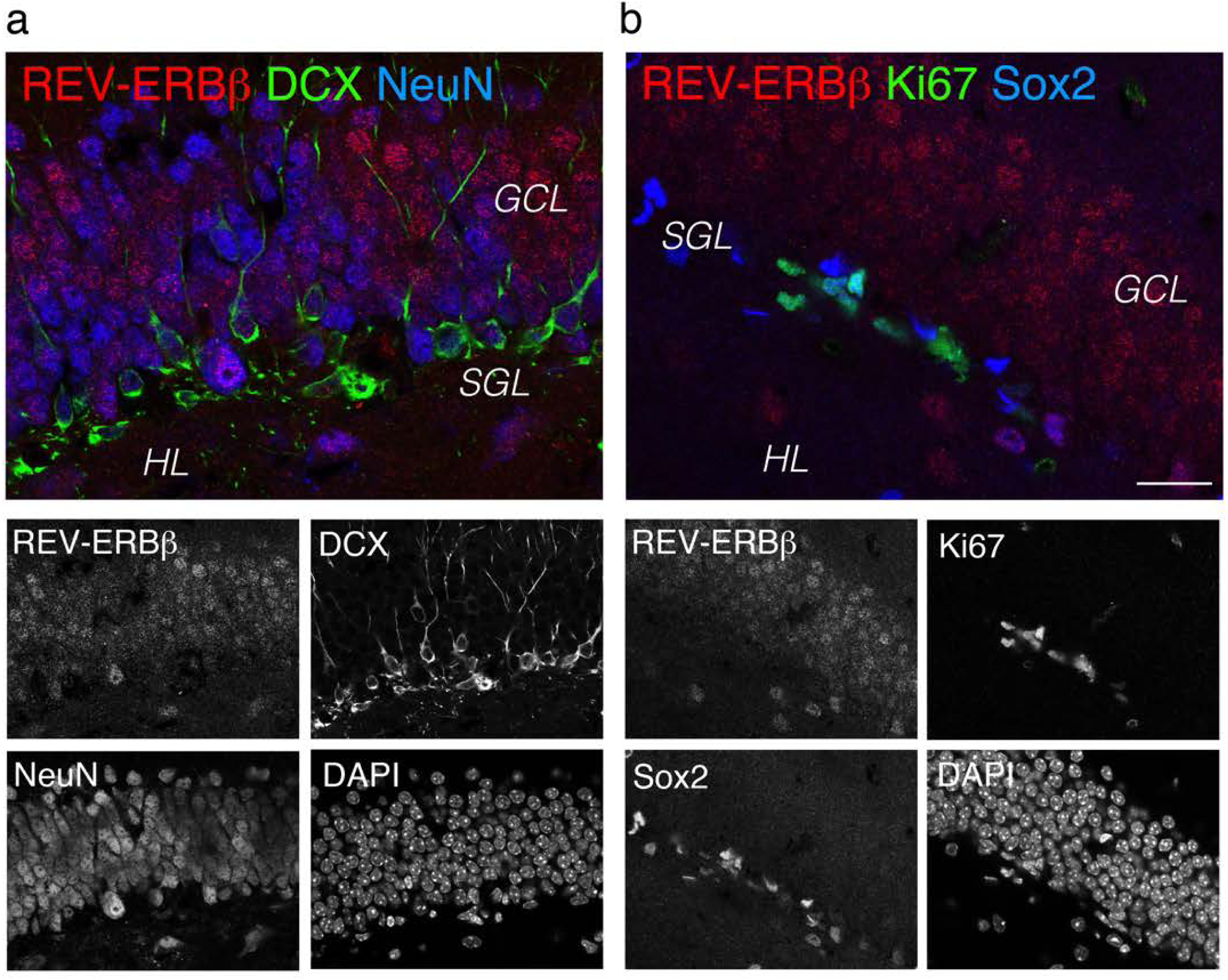
REV-ERBβ protein expression in the hippocampal dentate gyrus region of the adult mouse brain. The top main panel is a merged image of immunostained images of the hippocampal dentate gyrus of an adult mouse, and the bottom four sub-panels show images of each antibody and 4′ ,6-diamidino-2-phenylindole (DAPI) staining of the nucleus. **a.** Merged immunostaining image using anti-REV-ERBβ (red), doublecortin (DCX) (green), and neuronal nuclei (NeuN) (blue) antibodies (top) and single staining images with REV-ERBβ, DCX, and NeuN antibodies and DAPI (bottom). **b.** Merged image of immunostaining using anti-REV-ERBβ (red), Ki67 (green), and Sox2 (blue) antibodies (top) and single staining image of REV-ERBβ, Ki67, Sox2, and DAPI (bottom); *GCL*; granular cell layer, *SGL*; subgranular layer, *HL*; hilus, Scale bar represents 20 µm

### Expression of a dominant negative C-terminally deleted form of REV-ERBβ negatively affected dendrite formation in newborn neurons in the dentate gyrus of the adult mouse brain

To elucidate the function of REV-ERBβ in newborn neurons *in vivo*, a dominant negative form of *Rev-erbβ* was expressed in NSCs/NPCs of the dentate gyrus in the adult mouse brain. REV-ERBβ has a diversity domain (A/B region) in the nuclear hormone receptor in the N-terminal region and a DNA binding domain (C region) and a hinge region (D region) in the central region. A transcriptional repressor domain and ligand binding domain (E region) of the conserved domain structure are present in the carboxy-terminal region. Binding of this E region to the nuclear receptor co-repressor (NCoR) forms a transcriptional repression complex. A recombinant form of REV-ERBβ that lacked part of the D region and all of the E region was shown to inhibit the function of REV-ERBβ and act as a dominant negative form (Ramakrishnan et al. 2005). To clarify how this dominant negative form of REV-ERBβ acts on NSCs/NPCs in the hippocampal dentate gyrus of the adult mouse brain, truncated *Rev-erbβ* cDNA corresponding to amino acids 1–276, excluding the E region of REV-ERBβ, was incorporated into a retroviral expression vector. The green fluorescent protein (GFP) gene with an upstream internal ribosome entry site (IRES) was introduced downstream of the dominant negative form of *Rev-erbβ* under a CAG promoter, and infected cells became GFP positive.

This dominant negative *Rev-erbβ* expressing retroviral vector, together with VSV-G/gp plasmid DNA, was transfected into 293T cells, and the retroviral vector in the culture supernatant was collected, concentrated, and purified by centrifugation. The virus titer was adjusted to 1 × 10^7^ cfu ml^-1^ and injected into the hippocampal dentate gyrus of the adult mouse brain by stereotaxic microinjection. After 4 weeks, compressed maximum intensity projection (MIP) images of GFP-positive newborn neurons with a thickness of 25 µm were obtained by confocal laser microscopy, and a comparative analysis of single neurons from control samples and C-terminal-deficient dominant negative *Rev-erbβ*-expressing newborn neurons was performed (Fig. 2a). The dendritic length of GFP-positive, DCX-negative, and NeuN-positive newborn neurons was measured. Newborn neurons expressing the C-terminal deficient dominant negative form of *Rev-erbβ* had a shorter average dendritic length than controls (Fig. 2b; two-tailed unpaired *t*-test with Welch’s correction; df = 32.25; t = 5.485; *P* < 0.0001; n = 17 to 18). In addition, the average number of dendritic branches was lower in C-terminal deficient dominant negative *Rev-erbβ*-expressing newborn neurons than in control cells (Fig. 2c; two-tailed unpaired *t*-test with Welch’s correction; df = 32.85; t = 8.107; *P* < 0.0001; n = 17 to 18). These results indicated that expression of a dominant negative form of C-terminal-deficient *Rev-erbβ* suppresses part of the maturation process of newborn neurons during adult hippocampal neurogenesis.

**Fig. 2.**
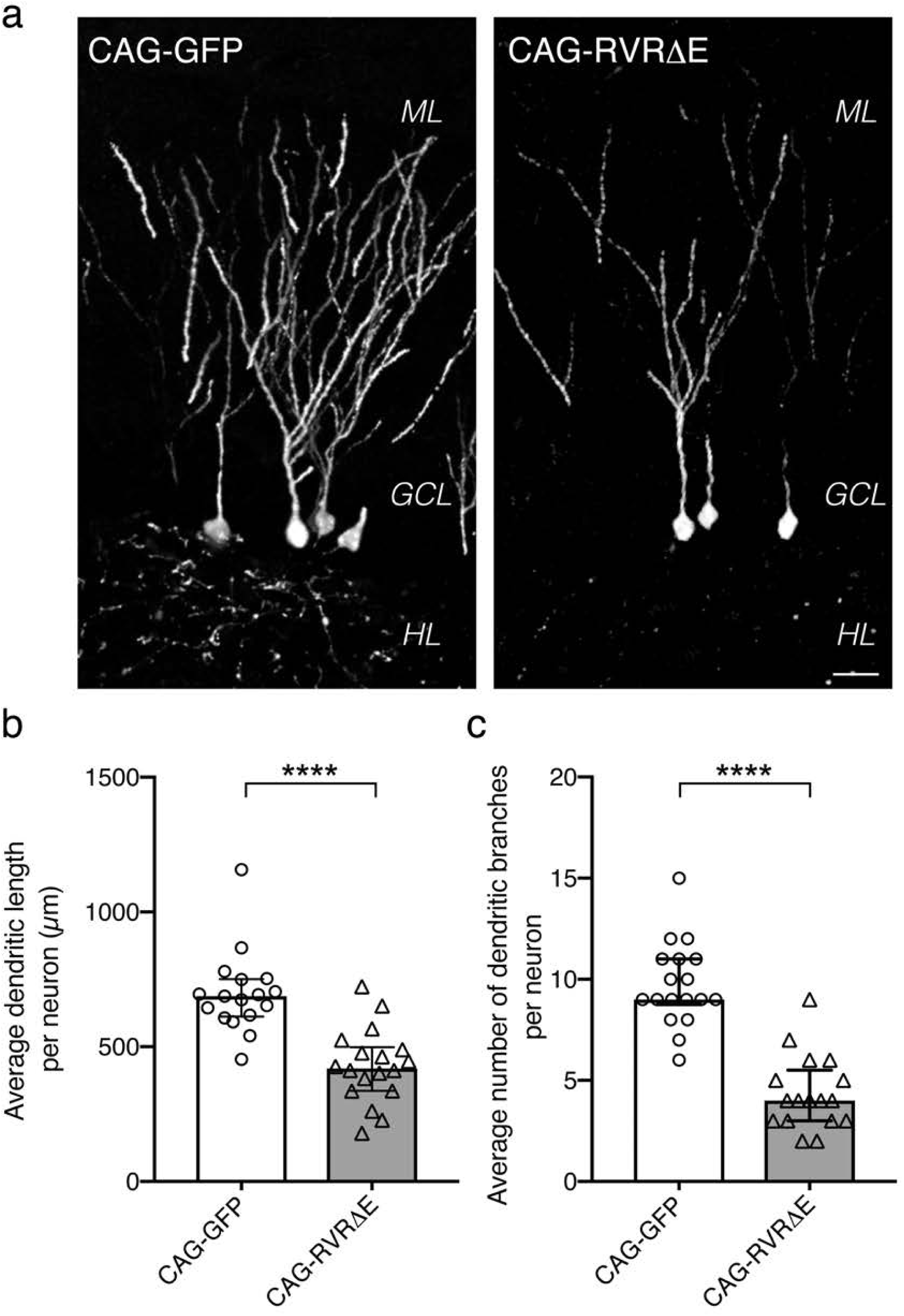
Comparative analysis of granule cell dendrites in the dentate gyrus of the adult mouse brain constitutively expressing *Rev-erbβ* with a C-terminal deletion. **a.** Maximum intensity projection images of newborn neurons in the hippocampal dentate gyrus of the adult mouse brain 4 weeks after infection with a control viral vector (CAG-GFP; right panel) or viral vector expressing *Rev-erbβ* with a C-terminal deletion (CAG-RVRΔE; left panel). *ML*, molecular layer; *GCL*, granule cell layer; *HL*, hilus. Scale bar represents 20 µm. **b, c.** Average dendritic length and average number of dendrite branches per new neuron in the hippocampal dentate gyrus of the adult mouse brain after 4 weeks of infection with a control viral vector (CAG-GFP, white bars, white circles) or a viral vector expressing *Rev-erbβ* with a C-terminal deletion (CAG-RVRΔE, gray bars, triangles), respectively. The average dendrite length and the average number of dendrite branches per new neuron in the dentate gyrus of the adult mouse brain after 4 weeks of infection were analyzed. Data are expressed as the mean ± s.e.m. or median with interquartile range (\*\*\*\**P* < 0.0001). Statistical comparisons were performed using an unpaired *t*-test with Welch’s correction

### Expression of a dominant negative form of C-terminally deleted *Rev-erbβ* affects spine formation in dendrites of newborn neurons during adult hippocampal neurogenesis

Next, the dendritic structure of newborn neurons expressing the C-terminal deleted dominant negative form of *Rev-erbβ* was examined. As shown in Supplemental data 1, MIP images of the dendrites of newborn neurons were obtained by confocal microscopy at high magnification, which showed that the dendrites of neurons expressing the dominant negative form of *Rev-erbβ* (CAG-RVRΔE) appeared to be less mature than those in controls (CAG-GFP). To obtain quantitative data on the detailed dendritic structure, a comparative analysis of the structure of the dendritic axis and the spines formed on the dendrites was performed by super-resolution microscopy using a structured illumination microscope (SIM) system, which performs structured illumination and provides higher resolution and more detailed image analysis. The SIM has a high degree of freedom in depth and pigment, and the spatial resolution is approximately twice that of confocal microscopy (approximately 100 nm). Using the SIM, a laser beam is projected onto the sample, and slit illumination is performed using a grating optimized for the excitation wavelength. The slit illumination can be moved in the phase direction as well as rotated to obtain more detailed information (Gustafsson 2005).

DCX-negative/NeuN-positive infected labeled newborn neurons were selected, and the dendritic details of GFP-positive newborn neurons were analyzed using a high magnification SIM system. Figure 3a shows a representative SIM image of a dendrite. The diameter of the dendritic axis was not different between control neurons and newborn neurons expressing the dominant negative form of *Rev-erbβ* (Fig. 3b; two-tailed unpaired *t*-test with Welch’s correction; df = 232.4; t = 0.7347; *P* = 0.4633; n = 124 to 131). Then, the dendritic spines of newborn neurons were examined. Spines are important synaptic connections in the construction of neural networks. However, the average dendritic spine length was not different between controls and newborn neurons expressing the dominant negative form of *Rev-erbβ* (Fig. 3c; two-tailed unpaired *t*-test with Welch’s correction; df = 239.6; t = 0.5064; *P* = 0.6130; n = 108 to 172). Interestingly, the diameter of the spine head was significantly smaller in newborn neurons expressing the dominant negative form of *Rev-erbβ* than in controls (Fig. 3d; two-tailed unpaired *t*-test with Welch’s correction; df = 262.5; t = 10.22; *P* < 0.0001; n = 107 to 159).

**Fig. 3.**
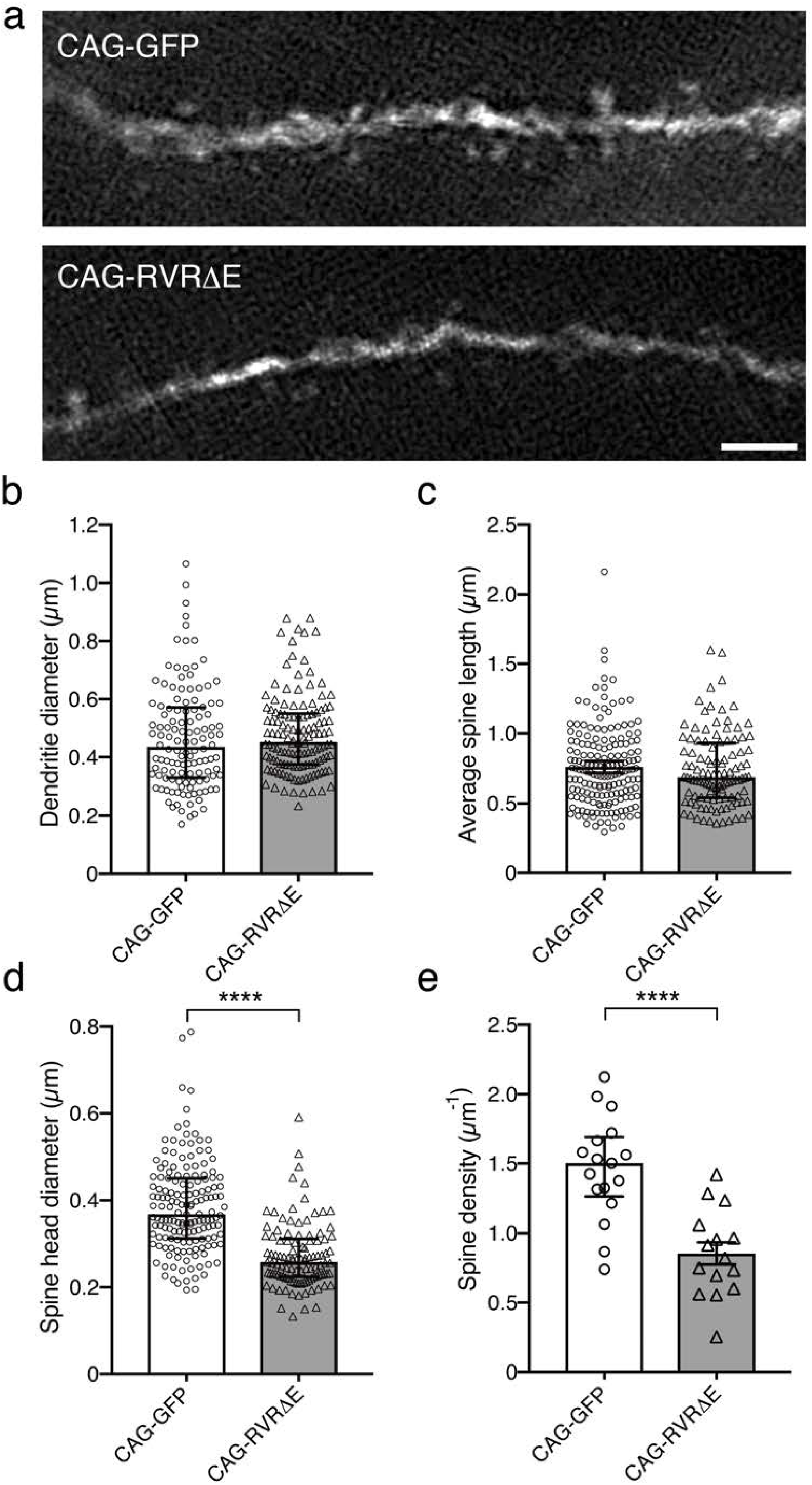
Comparative analysis of dendritic spine formation in newborn neurons in the dentate gyrus of the adult mouse brain expressing *Rev-erbβ* with a C-terminal deletion. The control vector virus (CAG-GFP) and the viral vector containing *Rev-erbβ* with a C-terminal deletion (CAG-RVRΔE) were introduced into neural stem/progenitor cells in the dentate gyrus of the adult mouse brain. Newborn neurons were analyzed by with a structured illumination microscope (SIM). **a**. Representative SIM images of dendrites of green fluorescent protein (GFP)-positive newborn neurons, Scale bar represents 2 µm. **b.** Diameters of dendritic axes of newborn neurons infected with the CAG-GFP (white bar) or CAG-RVRΔE (grey bar) retroviral vectors were measured from SIM images and plotted (CAG-GFP, white circles; CAG-RVRΔE, triangles). **c.** The spine lengths of newborn neurons infected with the CAG-GFP (white bar) or CAG-RVRΔE (grey bar) retroviral vectors were measured and plotted (CAG-GFP, white circles; CAG-RVRΔE, triangles). **d.** Comparative analysis of the diameter of spine heads of GFP-positive newborn neurons in SIM images. Spine heads of newborn neurons infected with CAG-GFP (white bar) and CAG-RVRΔE (grey bar) retroviral vectors were measured and plotted on a graph (CAG-GFP, white circles; CAG-RVRΔE, triangles). **e.** Comparative analysis of the spine density of GFP-positive newborn neurons in SIM images. The number of spines per micrometer of dendrite from randomly selected newborn neurons infected with the CAG-GFP (white bar) or CAG-RVRΔE (grey bar) retroviral vectors was counted and plotted (CAG-GFP, white circles; CAG-RVRΔE, triangles). The data are expressed as the mean ± s. e. m. or the median and interquartile range (\*\*\*\**P* < 0.0001). Statistical comparisons were performed using a two-tailed unpaired *t*-test with Welch’s correction

Furthermore, measurement of the density of spine bodies on the dendrite axis in control and dominant negative *Rev-erbβ*-expressing newborn neurons showed that expression of the dominant negative form of *Rev-erbβ* significantly reduced spine density (Fig. 3e; two-tailed unpaired *t*-test with Welch’s correction; df = 29.95; t = 5.054; *P* < 0.0001; n = 15 to 17). These results indicated that expression of the C-terminal deleted dominant negative form of *Rev-erbβ* does not affect the dendrite axial diameter or spine length during neurogenesis but has a negative effect on spine head formation and spine density.

### Expression of the dominant negative C-terminal deficient *Rev-erbβ* in the dentate gyrus of the adult mouse brain did not affect NSC/NPC proliferation and stage progression of neuronal differentiation of adult hippocampal newborn neurons *in vivo*

As shown in Figs. 2 and 3, expression of the dominant negative form of *Rev-erbβ* affected dendrite and spine formation during the transition from NSCs/NPCs to newborn neurons in the dentate gyrus of the adult mouse brain. The effect of the dominant negative form of *Rev-erbβ* on NSCs/NPCs was confirmed in the early stages of differentiation of newborn neurons. Figure 4a shows MIP images of GFP-positive cells obtained by confocal microscopy on day 4 after injection of the retroviral expression vector into the dentate gyrus of the adult mouse brain.

**Fig. 4.**
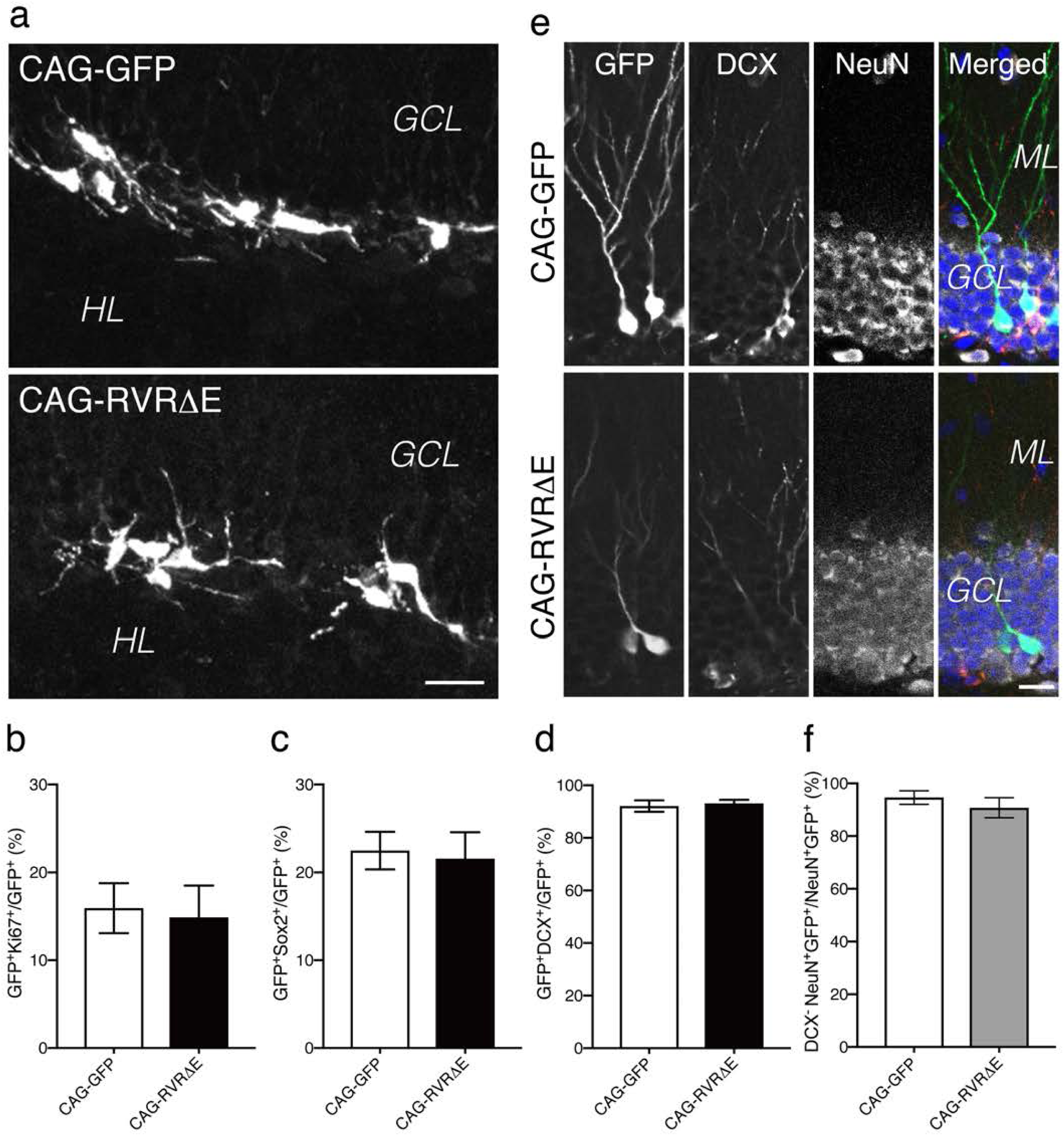
The effect of expression of *Rev-erbβ* with a C-terminal deletion on the proliferation and neuronal differentiation of neural stem/progenitor cells in the hippocampal dentate gyrus of the adult mouse brain. **a.** Representative maximum intensity projection images obtained by confocal microscopy of the hippocampal dentate gyrus of the adult mouse brain 4 days after infection with a control vector virus (CAG-GFP) or a viral vector expressing *Rev-erbβ* with a C-terminal deletion (CAG-RVRΔE). Scale bar represents 20 µm. **b.** Measurement of the proliferation rate of retroviral vector-infected cells after 4 days; the proliferation rate of CAG-GFP (white bar) or CAG-RVRΔE (black bar) infected cells was calculated by dividing the number of Ki67-positive (proliferation marker) green fluorescent protein (GFP)-expressing cells by the number of GFP-expressing cells. **c.** Measurement of Sox2-positive cells among virus vector-infected cells after 4 days; the number of Sox2-positive (neural stem/progenitor cell marker) cells among CAG-GFP (white bar) or CAG-RVRΔE (black bar) infected cells was counted. **d.** Measurement of doublecortin (DCX)-positive cells among virus vector-infected cells after 4 days; the number of DCX-positive (neuronal progenitor and immature neuron marker) cells among CAG-GFP (white bar) or CAG-RVRΔE (black bar) infected cells was counted. **e.** Representative immunostaining images of single sections obtained by confocal laser microscopy from the hippocampal dentate gyrus of the adult mouse brain after 4 weeks of infection with the control viral vector (CAG-GFP) or a viral vector expressing *Rev-erbβ* with a C-terminal deletion (CAG-RVRΔE). Merged images; GFP (green), DCX (red), neuronal nuclei (NeuN) (white), and 4′, 6-diamidino-2-phenylindole (DAPI) (blue). *ML*, molecular layer; *GCL*, granule cell layer. Scale bar represents 20 µm. **f.** Measurement of DCX-negative/NeuN-positive cells among viral vector infected cells after 4 weeks. The number of DCX-negative/NeuN-positive mature neurons among CAG-GFP (white bar) or CAG-RVRΔE (gray bar) infected cells was counted

CAG-GFP labeling in the top image indicates control cells. The cells exhibited morphology similar to type-1a and type-2a NSCs/NPCs and were interspersed between the HL and GCL. Similarly, cells infected with the dominant negative REV-ERBβ vector (CAG-RVRΔE) were also observed, but no morphological differences were observed between cells expressing the control CAG-GFP vector and the dominant negative CAG-RVRΔE vector (Fig. 4a). Because the viral vector used in this study is a retrovirus, the viral vector DNA is thought to mainly enter NSCs/NPCs during proliferation (Zhao et al. 2006). These results indicate that the dominant negative form of *Rev-erbβ* does not affect NPC morphology during proliferation. Next, the effect of dominant negative *Rev-erbβ* expression in NPCs on the proliferation mechanism was examined. Ki67 is a nuclear protein detected in the nucleus of proliferating cells. Sections of the hippocampal dentate gyrus region of adult mice infected with control virus and a viral vector expressing the dominant negative form of *Rev-erbβ* were immunostained with anti-Ki67 and anti-GFP antibodies. The number of cells positive for both anti-Ki67 and anti-GFP antibodies per observation area was detected by confocal fluorescence microscopy and divided by the number of GFP-positive cells. However, no significant difference was found in a comparison between control cells (CAG-GFP) and cells expressing the dominant negative form of *Rev-erbβ* (CAG-RVRΔE) (Fig. 4b; two-tailed unpaired *t*-test with Welch’s correction; df = 13.27; t = 0.2249; *P* = 0.8255; n = 8). To further investigate the stemness status, immunostaining was performed with an antibody against Sox2, an SRY box-containing transcription factor involved in the self-renewal mechanisms of NSCs and neural differentiation of NPCs that is also used as a marker for NSCs/NPCs. No significant difference was detected between control CAG-GFP and CAG-RVRΔE expressing cells regarding the positivity rates of anti-Sox2 and anti-GFP antibodies (Fig. 4c; two-tailed unpaired *t*-test with Welch’s correction; df = 12.62; t = 0.2489; *P* = 0.8075; n = 8). Immunostaining for DCX, which is expressed in NPCs and young progenitive neurons, showed no significant difference between control cells and cells expressing the dominant negative form of *Rev-erbβ* (Fig. 4d; two-tailed unpaired *t*-test with Welch’s correction; df = 9.971; t = 0.3881; *P* = 0.7061; n = 7). To examine the toxicity of the retroviral vector system, the dentate gyrus of adult mice was immunostained 4 days post-infection with an anti-active Caspase-3 antibody. The results showed no activated Caspase-3-positive apoptotic cells infected with either the control virus or the virus expressing the dominant negative form of *Rev-erbβ* (Supplemental data 2). These results suggested that in the hippocampal dentate gyrus of the adult brain, the dominant negative form of *Rev-erbβ*, lacking the C-terminal region, does not affect NSC/NPC proliferation, determination of the neuronal differentiation lineage of newborn neurons, or progression of the differentiation stage. Next, to elucidate the influence of the dominant negative form of *Rev-erbβ* on neuronal fate determination and differentiation processes, NSCs/NPCs in the adult hippocampal dentate gyrus were infected with CAG-RVRΔE, and immunostaining for neural differentiation markers was performed after 4 weeks. Newborn neurons produced from NSCs/NPCs form DCX-positive neuronal progenitor cells and finally differentiate into DCX-negative/NeuN-positive neurons. Single-section analysis of confocal laser microscopy images showed that control newborn neurons were negative for the neuronal progenitor cell marker DCX and positive for NeuN, a marker of differentiated neurons. However, newborn neurons expressing the C-terminal deleted dominant negative form of *Rev-erbβ* were also negative for DCX and positive for NeuN (Fig. 4e). Among GFP-positive newborn neurons, DCX-negative/NeuN-positive cells were counted and compared between control cells and cells expressing the dominant negative *Rev-erbβ*. No significant difference was detected between the two groups (Fig. 4f; two-tailed unpaired *t*-test with Welch’s correction; df = 10.54; t = 0.8394; *P* = 0.4199; n = 8). Therefore, NSCs/NPCs underwent progressive neurogenesis, regardless of whether they expressed the dominant negative form of *Rev-erbβ*. Taken together, these results suggested that the dominant negative form of *Rev-erbβ* lacking the C-terminus does not interfere with the normal differentiation and maturation process of newborn neurons in the hippocampal dentate gyrus of the adult mouse brain.

## Discussion

In this study, the expression and localization of the REV-ERBβ protein was first investigated, focusing on the hippocampal dentate gyrus of the adult mouse brain. Analysis of REV-ERBβ protein expression in cultured mouse NSCs showed localization in the nucleus and cytoplasm (Shimozaki 2018). However, the *in vivo* results in the hippocampal dentate gyrus showed that most REV-ERBβ-positive cells were NeuN-positive neurons, including a small number of cells that were also positive for DCX. No REV-ERBβ protein expression was found in proliferating NSCs/NPCs, including Sox2-positive and Ki67-positive cells. This discrepancy may be caused by optimization of gene expression. That is, the *Rev-erbβ* gene may be switched on in Sox2-positive and REV-ERBβ-negative cells during the establishment of an *in vitro* NSC culture because cultured NSCs are in a constant state of proliferative signaling activated by FGF or EGF to ensure that the stemness gene is permanently expressed (Zhang et al. 2008). Another possibility is that cultured NSCs may originate from unique Sox2-positive cells because a small number of Ki67-negative/Sox2-positive cells were also observed to express REV-ERBβ (Fig. 1b). Interestingly, a ligand of REV-ERBβ, SR9009, has been reported to inhibit glioma cell growth and to act as a cancer treatment drug (Sulli et al. 2018; Wagner et al. 2019; Dong et al. 2019). In this study, REV-ERBβ was mainly expressed in differentiated neurons, although a small number of cells expressing REV-ERBβ was detected among Ki67-negative/Sox2-positive cells. Though the details of how cancer stem cells (CSCs) are produced are still unclear, CSCs are known to share some properties with tissue stem cells including NSCs. CSCs are responsible for tumor recurrence after chemotherapy. Using the promoter activity of the NSC marker gene Nestin as an indicator, CSCs were identified and targeted to reduce tumor recurrence in mice, which increased their survival (Chen et al. 2012). Therefore, regulation of *Rev-erbβ* gene expression may be an important aspect of future neurogenesis and CSC research, as it could provide insights into the mechanisms by which SR9009 exerts anti-tumor activity against gliomas.

The retroviral vector used for gene expression in this study expresses an inserted gene and an enhanced GFP gene located downstream of the IRES sequence. The constitutive expression of the inserted gene in proliferating cells and cell labeling with GFP were carried out simultaneously, allowing *in vivo* cell lineage and cell biological analysis. Inoculation of this vector system into the SGL of the hippocampal dentate gyrus of the adult mouse brain comprises a unidirectional *in vivo* gene expression system. A small number of proliferating type-2a NSCs and type-2b NPCs were infected, and after 4 weeks, approximately 70% of cells were newborn neurons (Zhao et al. 2006). Using this system, expression of *Ascl1* in NSCs/NPCs can induce cell lineage conversion to oligodendrocytes (Jessberger et al. 2008), while expression of the transcription factors Prx1/PRRX1 can also induce de-differentiation to type-1 NSCs and oligodendrocyte progenitor-like cells (Shimozaki et al. 2013). The C-terminus deletion mutant of REV-ERBβ used in this study has been reported to decrease genes involved in fatty acid and lipid absorption and regulators of muscle formation while increasing the expression of pro-inflammatory cytokines when expressed in mouse myogenic C2C12 cells (Ramakrishnan et al. 2005). However, expression of this mutant in NSCs/NPCs of the hippocampal dentate gyrus had no effect on cell lineage, proliferation, or cell death (Fig. 4).

Because the infected cells were mainly proliferative type-2a and -2b cells, it was expected that if resistance to differentiation or maintenance of undifferentiated cells was induced, the cells would remain type-2a NSCs or return to type-1 NSCs. In the present study, however, neural progenitor differentiation in *Rev-erbβ*-dominant negative expressing cells proceeded at a similar rate to that in controls. This suggests that the action of REV-ERBβ varies among cell types and is flexibly regulated by context-dependent mechanisms of intracellular molecules. However, this does not indicate that REV-ERBβ itself is irrelevant in regulating proliferation and cell lineage. Although the system used in this study was designed to produce a dominant negative effect, there are often differences in the results obtained between various experimental methods used to inhibit the function of a target factor. Gene knockdown and gene knockout can also change the phenotype of cells because of their differential effects on target genes (El-Brolosy et al. 2019). Therefore, to more accurately understand the *in vivo* function of REV-ERBβ, further analyses using a wide range of methods, including gene knockdown and gene knockout methods, will be required to reach a more comprehensive conclusion.

C-terminal deletion of the *Rev-erbβ* gene and expression as a dominant negative form in NSCs/NPCs of the hippocampal dentate gyrus of the adult mouse brain reduced the dendrite outgrowth and branching ability of newborn neurons (Fig. 2). This suggests that REV-ERBβ affects dendrite growth in newborn neurons. Chronic stress has also been implicated in inhibiting the proliferation and maturation mechanisms of NSCs in the hippocampal dentate gyrus, and depressed patients show reduced hippocampal size and function (MacQueen et al. 2003; Schoenfeld et al. 2017; Numakawa et al. 2017; Belleau et al. 2019).

Neurogenesis is involved in antidepressant action, and the antidepressant fluoxetine has also been reported to increase the number of newborn neurons (Malberg et al. 2000) and promote the development and maturation of dendrites in newborn neurons (Amellem et al. 2017). Fluoxetine inhibits serotonin reuptake, thereby increasing the serotonin concentration between synapses.

Serotonin integrates the neural activity and circadian rhythms associated with sleep and wakefulness deep within the brain (Miyamoto et al. 2012). Although the direct causal relationship between the circadian rhythm regulator REV-ERBβ and depression is not known, it will be an interesting challenge to analyze the potential of REV-ERBβ as a target for antidepressant signaling. The present study also suggested that REV-ERBβ regulates dendritic spine size and density in addition to dendrite growth (Fig. 3). Recently, REV-ERBβ knockout mice were shown to exhibit anxious behavior and obsessive-compulsive disorder-like symptoms (Banerjee et al. 2014). According to Banerjee et al. stimulation with multiple REV-ERB agonists ameliorates these symptoms. Moreover, my recent results showed that an REV-ERB agonist, SR9009, regulates neurites in cultured NSCs from adult rats in a concentration-dependent manner. SR9009 induces neurite outgrowth at low concentrations and inhibits neurite outgrowth at high concentrations via REV-ERBβ during neuronal differentiation from cultured NSCs (Shimozaki 2021). Modulation of REV-ERBβ activity would allow for regulatory control of dendrites and spines in newborn neurons *in vivo*, but this issue remains to be investigated. Furthermore, one cause of Alzheimer’s disease (Gu et al. 2020) is cytoplasmic aggregation of the transactive response DNA-binding protein of 43 kDa (TDP-43) in neurons.

In Alzheimer’s disease, CK1ε expression is upregulated, suggesting that CK1ε is involved in the aggregation of TDP-43 by increasing its phosphorylation level. However, phosphorylation of the N-terminus of REV-ERBβ by CK1ε enhances its localization to the cytoplasm. Because the REV-ERBβ protein is expressed in the nucleus of newborn neurons (Fig. 1), the enhanced cytoplasmic localization of REV-ERBβ may affect dendrite formation and synaptogenesis. The REV-ERB agonist SR9009 has also been reported to upregulate PSD-95 expression and increase the number of synapses in the hippocampus of mouse models of Alzheimer’s disease (Roby et al. 2019). Therefore, the relationship between REV-ERBβ and Alzheimer’s disease is also considered to be important.

Neurogenesis in the adult brain is only one part of the *in vivo* role of REV-ERBβ. In addition to circadian rhythm control, REV-ERBs play an important role in cholesterol metabolism (Sitaula et al. 2017). They are also involved in muscle formation and muscular dystrophy (Mayeuf-Louchart et al. 2017; Welch et al. 2017) and autoimmune disease (Amir et al. 2018; Ishimaru et al. 2019; Chang et al. 2019), and an association between heart failure and myocardial infarction (Zhang et al. 2017; Stujanna et al. 2017; Reitz et al. 2019) has also been found. The exact role of REV-ERBβ *in vivo* will be further elucidated by using mice genetically modified with the Cre/loxP system, combining tissue-specific and differentiation stage-specific gene promoters, and administering REV-ERB agonists to individual mice. *In vivo* cell-specific analysis of the dominant negative form of REV-ERBβ in this study revealed a new role for REV-ERBβ at the single cell level. The results of this study will provide important scientific insights into adult brain neurogenesis and psychiatric disorders.

## Materials and methods

### Construction of a dominant negative form of *Rev-erbβ* and insertion into a retroviral expression vector

DNA fragments of C-terminally deleted *Rev-erbβ* (1–829 bp) were obtained by PCR amplification of cDNA prepared from mRNA extracted from cultured mouse NSCs (Zhang et al. 2008; Shimozaki 2017, 2018). The PCR-amplified DNA fragment (RVRΔE) was TA-cloned, sequenced by the Sanger method, and incorporated into the retroviral expression vector pCAG-IRES-EGFP (Zhao et al. 2006; Jessberger et al. 2008). This vector drives the gene via the cytomegalovirus enhancer and chicken β-actin promoter but also expresses enhanced GFP using the IRES sequence. A retrovirus was generated from this vector plasmid as previously described (Zhao et al. 2006). The titers of the centrifuged viruses ranged from 1.2 to 9 × 10^7^ cfu ml^-1^ and were finally adjusted to 1 × 10^7^ cfu ml^-1^.

### Injection of retroviral vectors into the hippocampal dentate gyrus of the adult mouse brain and immunostaining analysis

Animal procedures were carried out according to protocols approved by the Animal Experimentation Committee of Nagasaki University. For this study, 8 to 10-week-old female C57Bl/6JJmsSlc mice purchased from SLC Japan were used. Mice were injected with 1 µl of the virus containing CAG-GFP (pCAG-IRES-EGFP) or CAG-RVRΔE (pCAG-RVRΔE-IRES-GFP) in the left or right portion of the dentate gyrus of the same individual using a stereotaxic injector (coordinates from bregma: −2 mm anterior/posterior, ±1.5 mm medial/lateral and −2.3 mm dorsal/ventral). Virus vector-inoculated mice were perfusion-fixed with 0.9% NaCl and 4% paraformaldehyde, and then the whole brain was placed in 30% sucrose solution at 4°C for 2 days. Sections with a thickness of 40 µm were prepared from the fixed whole brain using a microtome. The sections were then treated with a blocking solution consisting of Tris buffered saline (TBS) solution and 5% donkey serum and kept at room temperature for 2 hours. Then, the primary antibody was added, and the sections were incubated at 4°C overnight. On the following day, the sections were washed with TBS solution, and then the secondary antibody was added. The reaction was carried out at room temperature for 2 hours. The primary antibodies included anti-GFP (Aves Labs), anti-NR1D2 (BioLegend), anti-DCX (Santa Cruz), anti-NeuN (Cell Signaling), anti-Sox2 (Cell Signaling), anti-Ki67 (Abcam), and anti-active Caspase-3 (BD Pharmingen). Anti-rat, goat, and rabbit DyLight™ 488, AX549, and AX649 IgG (Jackson ImmunoResearch Laboratories) were used as secondary antibodies as appropriate. Images of immunostained sections were obtained using an LSM800 confocal laser microscope (ZEISS). Representative images are shown as single section images or compressed images with 25 µm MIP. To assess the dendrite length and number of dendritic branches after 4 weeks, 17–18 immunostained GFP-positive cells were randomly selected from brain sections obtained from four to six virus-injected mice and measured using ZEN3.2 blue edition software (ZEISS). The percentages of GFP-, Ki67-, Sox2-, DCX-, and NeuN-positive cells were calculated from Z-stack images (25 µm) of two to four brain sections per slide for each protein and averaged for seven to eight slides from six to nine virus-injected mice to determine the phenotype of virus-labeled cells at 4 or 28 days after injection. Dendritic fine structure and spine analyses were performed with a SIM (ELYRA PS.1/ LSM710 microscope, ZEISS). Among the randomly selected GFP-positive cells, the width of 124–131 dendrites, length of 108–172 spines, size of 107–159 spine heads, and number of spines in 15– 17 areas were measured using the application software described above.

## Statistical procedures

All data were statistically evaluated using Prism 7 statistical software (GraphPad). An unpaired *t*-test with Welch’s correction was performed, and the data are presented graphically as the mean and standard error of the mean (± s. e. m.) or median and interquartile range. *P*-values less than 0.05 were considered statistically significant.

## Acknowledgments and Declarations

I thank Lisa Kreiner, PhD, from Edanz (https://www.jp.edanz.com/ac) for editing a draft of this manuscript and helping to draft the abstract.

## Funding

This work was supported by JSPS KAKENHI Grant Number JP23K10962 and by a research grant from the Nagasaki Medical Alumni Association.

## Data Availability

The data that support the findings of this study are available from the corresponding author, K. Shimozaki, upon reasonable request.

## Conflict of interest

The author has no conflicts of interest directly relevant to the content of this article.

---

**Supplemental data 1.**
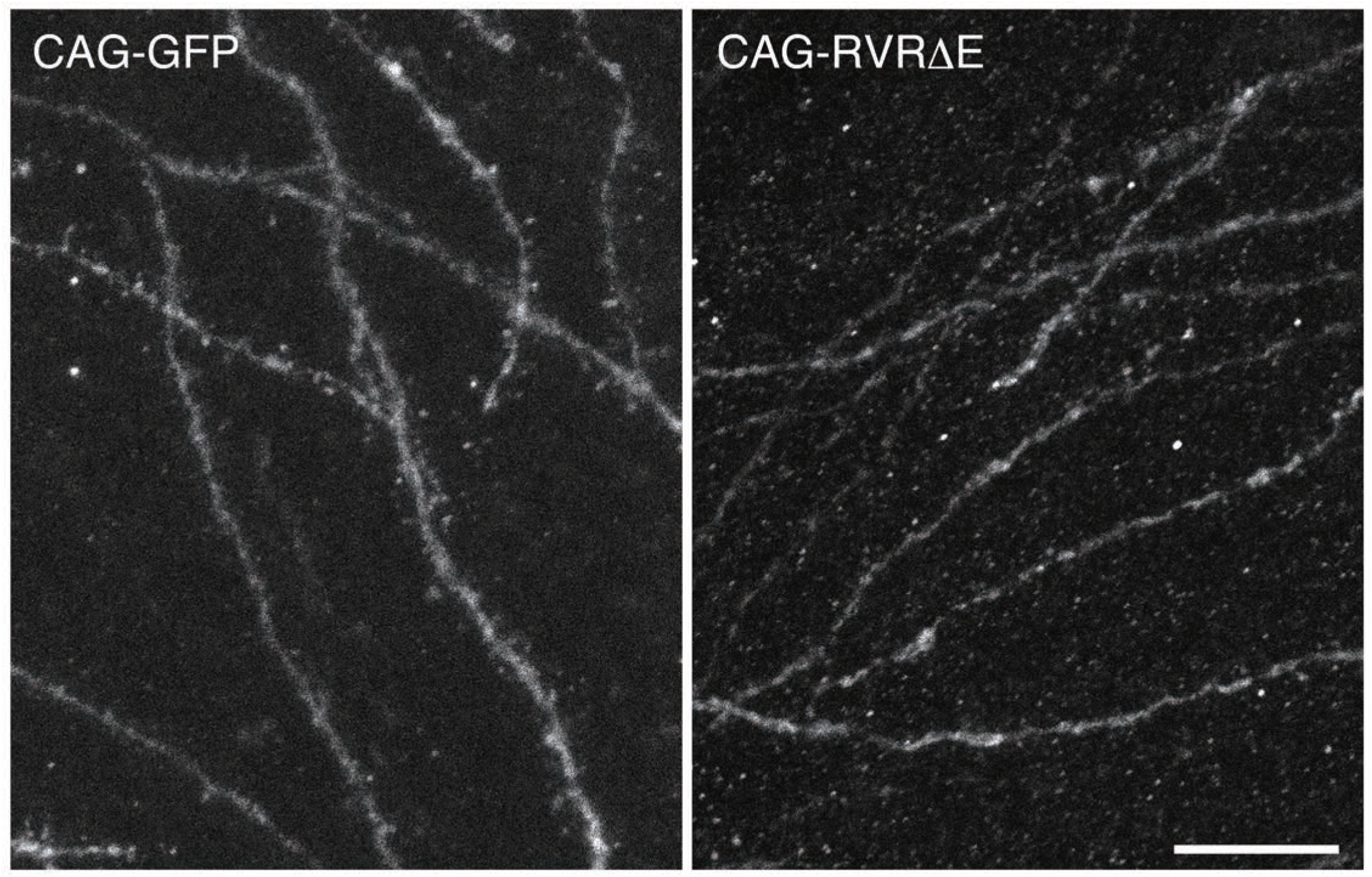
Maximum intensity projection images (thickness 25 µm) of dendrites of newborn neurons in the hippocampal dentate gyrus of the adult mouse brain at 4 weeks after infection with a control viral vector (CAG-GFP; right panel) or a viral vector expressing *Rev-erbβ* with a C-terminal deletion (CAG-RVRΔE; left panel). Scale bar represents 10 µm

**Supplemental data 2.**
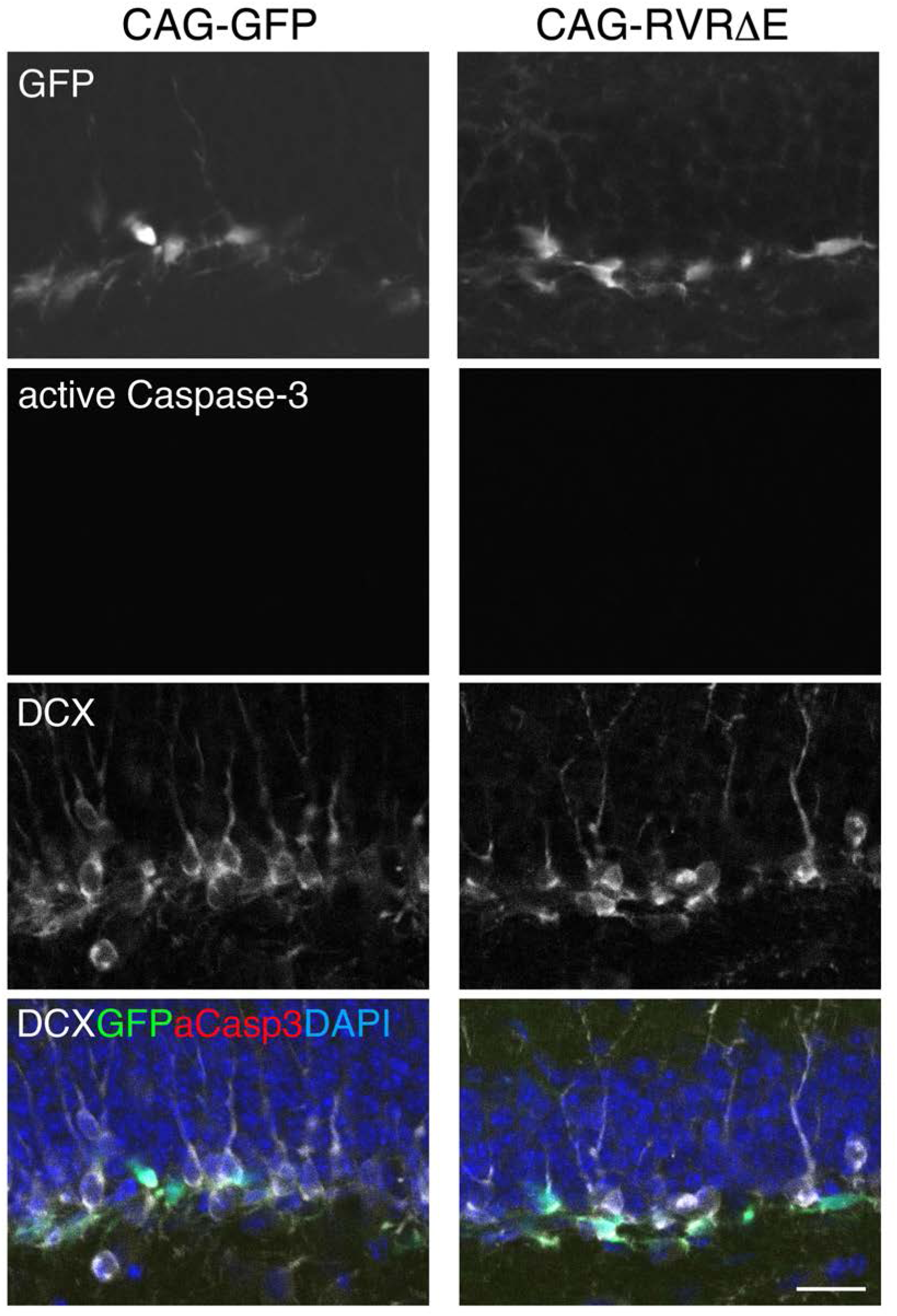
Lack of apoptotic activity of C-terminal deleted REV-ERBβ in the dentate gyrus of the adult mouse brain. The control viral vector (CAG-GFP) and a viral vector expressing *Rev-erbβ* with a C-terminal deletion (CAG-RVRΔE) were infected into neural stem/progenitor cells in the dentate gyrus of the adult mouse brain. After 4 weeks, newborn neurons were immunostained with an anti-active Caspase-3 antibody, and images of single sections obtained by confocal laser microscopy were analyzed. Merged images; green fluorescent protein (GFP) (green), doublecortin (DCX) (white), active Caspase-3 (aCasp3) (red), 4′ ,6-diamidino-2-phenylindole (DAPI) (blue). Scale bar represents 20 µm

## Notes

### Competing Interest Statement

The authors have declared no competing interest.

